# Moving in complex environments: a biomechanical analysis of locomotion on inclined and narrow substrates

**DOI:** 10.1101/382028

**Authors:** Christofer J. Clemente, Taylor J.M. Dick, Rebecca Wheatley, Joshua Gaschk, Ami Fadhillah Amir Abdul Nasir, Skye F. Cameron, Robbie S. Wilson

## Abstract

Characterisation of an organisms’ performance in different habitats provides insight into the conditions that allow it to survive and reproduce. In recent years, Northern quolls (*Dasyurus hallucatus*)—a medium-sized semi-arboreal marsupial native to northern Australia—have undergone significant population declines within open forest, woodland and riparian habitats, but less so in rocky areas. To explore this decline, we quantified the biomechanical performance of wild Northern quolls over inclined narrow (13 mm pole) and inclined wide (90 mm platform) substrates. We predicted that quolls may possess biomechanical adaptations to increase stability on narrow surfaces, which are more common in rocky habitats. Our results display that quolls have some biomechanical characteristics consistent with a stability advantage on narrow surfaces. This includes the coupled use of limb pairs and high grip torques (Max = 68.9 N.m, Min = −69.9 N.m), highlighting their ability to generate corrective forces to counteract the toppling moments commonly encountered during gait on narrow surfaces. However, unlike other arboreal specialists such as primates, speed was constrained on narrow surfaces, and quolls did not adopt diagonal sequence gaits. Quolls appear to adopt similar performance characteristics to cats and dogs which may limit their ability to outperform these key predators (invasive or otherwise) in particular habitats. This overlap may contribute to the declining population of Northern quolls on mainland Australia.

## Introduction

An ecological niche is the set of environmental conditions that enable a species to grow and reproduce (Schoener, 2009). The fitness of an organism is largely defined by its performance in any environment which in turn depends upon morphology and physiology (Garland Jr and Losos, 1994, Arnold, 1983). Thus, characterisation of a species’ performance across different habitats provides insight into the conditions that allow for an animal to survive and reproduce. All behaviours require movement, thus the niche that an animal occupies is largely dependent on movement capabilities; for example, if an animal cannot climb trees it cannot fill a niche that is wholly or partially arboreal. Further, an animal’s performance will change with respect to environmental conditions that hinder movement. Understanding the locomotor capabilities of a species is key for understanding its niche, which may allow us to quantify the habitat requirements for conservation. One species of conservation significance in Australia is the Northern quoll (*Dasyurus hallucatus*; Gould, 1842).

Northern quolls are primarily nocturnal, partially arboreal marsupial carnivores, found in grassy or rocky habitats across northern Australia (Schmitt et al., 1989, Oakwood, 2000). The Northern quoll has declined from a broad distribution across northern Australia to several disjunct populations often centred on rocky plateaus with local extinctions primarily occurring in lowland savanna (Braithwaite and Griffiths, 1994, Morris, 1996, Braithwaite and Muller, 1997). Dingoes and feral cats are the main predators of Northern quolls, and both have been historically present across their range for 4000 and over 100 years, respectively (Burbidge et al., 1988, Corbett, 1995). The reason for the recent rapid decline in population is therefore complex. However, it has been hypothesised that habitat loss or fragmentation (due to land clearing, altered fire regimes, or grazing by invasive herbivores) can leave smaller mammalian species more vulnerable to predation (Newsome, 1975, Burbidge and McKenzie, 1989, McKenzie et al., 2007). In a study on quoll survival, Oakwood (2000) reported that all Northern quolls killed by predators were killed within forest, woodland and riparian habitats, and that few kills occurred in rocky outcrops. Also, females whose home ranges included a greater proportion of rocky habitat, were more likely to survive to a second breeding season. This suggests that rocky habitats may increase the survival of Northern quolls.

Greater survival in rocky habitats could be due to either a higher density of refugia or a lower abundance of large predators in these habitats. Dingoes, a major predator of quolls, mainly hunt on floodplains during the dry season and in the forest during the wet season, and are most successful in open habitats (Corbett, 1995). Dingoes were never observed on the rocky hills during this study, suggesting that the locomotion of dingos may be compromised in these habitats relative to open areas. Similar strategies have been shown in feral cats, which occupy open habitats more frequently than complex ones (Hohnen et al., 2016) and display better hunting success in these open habitats when compared to complex ones (McGregor et al., 2015).

The differential survival of a species across various habitats will be due to changes in their detection by predators or ease of capture after detection. Studying the performance of a species vulnerable to predation across various types of terrain allows one to make predictions about the relative performances between predator and prey. This can provide the basis for a mechanistic understanding of predation and its conservation significance.

Our study examined the kinematics and kinetics of Northern quolls in both simplified terrestrial and arboreal environments, designed to represent aspects of their natural habitat. We categorised their biomechanics across two different substrates: an inclined wide platform, and an inclined narrow pole. We then compared the quolls’ biomechanics on each of these substrates with biomechanical strategies used by arboreal specialists (such as primates and possums) and terrestrial specialists (such as rats, cats, and dogs) to compare the relative performances of quolls to these groups of animals. We hypothesized that within complex arboreal-like habitats quolls would modify biomechanical characteristics to be more reflective of arboreal species, and therefore help to explain their increased survival in these habitats.

## Materials and Methods

### Morphology

Northern quolls were trapped on Groote Eylandt, Northern Territory, Australia, between July and August 2013 using Tomahawk original series cage traps (20 × 20 × 60 cm; Tomahawk ID-103, Hazelhurst, Winconsin, USA) baited with canned dog food. Traps were set overnight and checked early in the morning (no later than 7:30 am) to avoid quolls being subjected to warmer parts of the day. A total of 24 individual quolls (9 males, 15 females) were captured throughout the study period. Each quoll was taken to the Anindilyakwa Land and Sea Ranger Research Station for subsequent experiments. All individuals received a micro-chip (Trovan nano-transponder ID-100, Keysborough, Australia) that was placed subcutaneously between the shoulder blades to ensure identification during any subsequent recaptures. Research methodologies were approved by the University of Queensland animal ethics committee (SBS/541/12/ANINDILYAKWA/MYBRAINSC) and were conducted under the Northern Territory Parks and Wildlife Commission (permit number: 47603).

Body mass was measured for each individual using an electronic balance (±0.1 g; A & D Company Limited HL200i, Brisbane, Australia) and eleven morphological variables were each measured three times using digital calipers, with an average used as the overall measure (Whitworth, Brisbane, Australia, ±0.01 mm) (Wynn et al., 2015). These body dimensions were included as covariates in our initial analysis, but none appeared to have a significant effect on our results and thus were not included in further analysis. Mean body mass was 384.3 ± 21.5 g, head width (widest point of jaw) 36.07 ± 0.47 mm, head length (from nuchal crest to tip of snout) 67.22 ± 0.73 mm, body length (nuchal crest to base of tail) 179.92 ± 3.32 mm, right and left fore-limb length (radius-ulna) 50.91 ± 0.75 mm, right and left hind-limb length (tibia-fibula) 63.75 ± 0.79 mm, right and left hind-foot length (heel to claw base) 38.49 ± 0.48 mm, tail width (maximum tail diameter) 14.54 ± 0.43 mm and tail length (base to tip of tail) 228.48 ± 4.49 mm. To minimize long-term stress on the animals, all performance measures and tests were completed within 6 hours of capture, after which the animals were released at the exact point of capture.

### Kinematics and kinetics

To assess the biomechanics of Northern quolls, we filmed each individual as it moved over two different raised platform trackways. Trackways were suspended within a Perspex box (2600 × 470 × 160 mm) with a release and escape box at the start and end, respectively (Figure 1).

**Figure 1.**
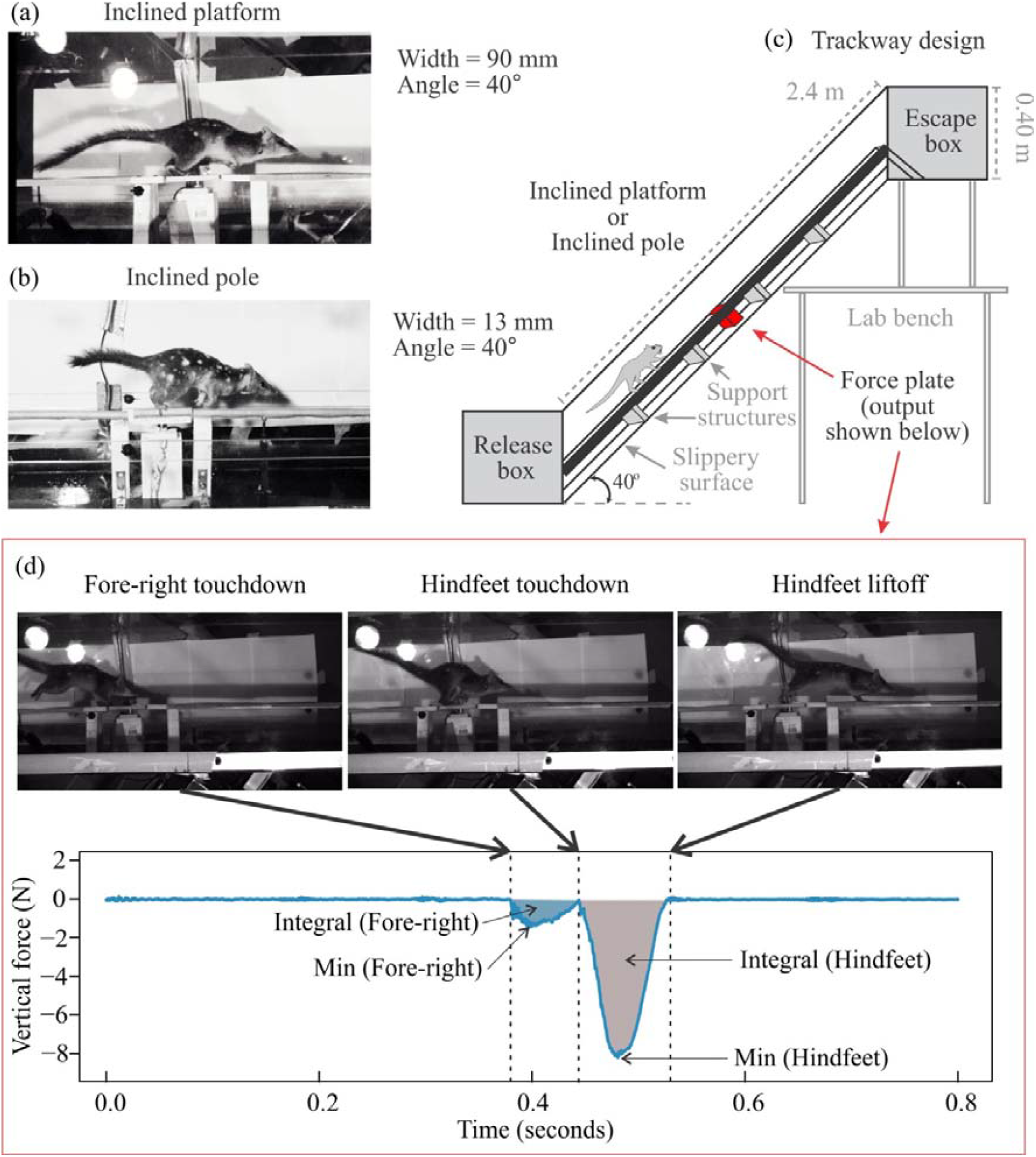
Schematic of experiment setup: a) Northern quoll (*Dasyurus hallucatus*) running on inclined platform, b) quoll running on inclined narrow poll, c) schematic of custom made experimental setup showing location of force plate, and d) vertical substrate reaction force trace over time for a single forelimb followed by both hindlimbs with representative images shown above. Shaded regions represent integral with respect to time (impulse) for the fore-right (blue) and hind-left and hind-right combined (grey).

Quolls were run along the two trackways (angle, pole) in a random order, to represent different habitat types that a quoll is likely to encounter in its natural environment. The inclined platform was a 90 mm wide plank of wood, covered in fine sandpaper (P120) to provide traction at a 40° angle, and represented a terrestrial environment. The inclined pole was a 12 mm diameter wooden dowel running the length of the box at a 40° angle, and represented an arboreal environment. A third flat platform trackway (90 mm wide platform) was also tested, but later excluded (see excluded strides). Both tracks ran the length of the box with a force platform (Nano-17 titanium, ATI instruments) placed level and centrally with the trackway, with either a 90 × 90 mm platform or a 90 mm long doweling attached to its top.

Two cameras (Fastec IL-3, Fastec Imaging, 1280 × 700 pixels, @ 250 fps) were positioned approximately perpendicular to each other, to capture a ventral/lateral, and lateral/dorsal view. These camera positions were used to ensure footfalls were visible in both camera views, yet allowed enough variation in viewing angle that 3-dimensional position information was still accurate. Cameras were synchronized using internal triggers. The cameras were calibrated using a wand based camera calibrate, implemented in the Argus software in Python Ver 2.1, (Jackson et al., 2016). The Matlab GUI DLTdv5 (Hedrick, 2008) was then used to track individual feet and a single spot on the back, lose to the body centre of mass.

### Analysis

#### (i) Timing parameters and gait

Feet were digitised for the duration of contact with the surface of calibrated racetrack. A mark near the body centre of mass was used to calculate speed. Footfall sequence and gait parameters were calculated using antero-posterior sequence methods (Abourachid, 2003, Chadwell and Young, 2015). The first forefeet touchdown defined the start of a sequence and its subsequent touchdown defined the end of the sequence. F-lag, H-lag and P-lag were calculated as the temporal lag between the mid-stances of the two forefeet, hindfeet and averaged across the ipsilateral feet respectively. We used the gait parameters described by Abourachid (2003) and Chadwell and Young (2015). Speed was calculated for the duration of the sequence, and duty factor was calculated as the average duty factor of feet within a sequence. We examined the effects of, and interaction between, speed and surface for the time lag between footfalls of the forefeet (F.lag), hindfeet (H.lag) and the difference between the fore and hindfeet (P.lag) to determine which pairs of limbs may be contributing most to increasing stability on narrow perches.

#### (ii) Spatial parameters

We further examined the effects of the absolute distance between the left and right fore-foot (F.dist), the left and right hindfoot (H.dist) and the mean distance between the forefoot pair and the hindfoot pair (P.dist). Since the lateral distance effects are likely to be constrained on narrow supports we further examined the distance in the fore-aft axis for each variable (F.distX, H.distX and P.DistX respectively).

#### (iii) Substrate reaction forces

Substrate reaction forces were recorded for each foot individually or for multiple feet simultaneously using an ATI Nano17 6 dof micro force transducer (ATI instruments). The forceplate was calibrated by the company with estimated error of < 0.75% in the x, y, and z axes. The force plate was aligned with the x-axis representing the fore-aft direction, y representing the lateral direction, and z representing vertical forces. The transducer output forces in these three axes along with torques about each axis. Torques about the axes are likely dependent on the position of the footfall relative to the centre of the force transducer and therefore of limited use, with the exception of the torque about the x-axis on the narrow trackway. In this setup, the narrow pole was less than the width of the foot. Thus, we assume the position of the footfall must more or less align with the x-axis of the force plate, and therefore we included this torque in our analysis.

Strides where any foot partially stepped on the plate were excluded from further analysis. Forces were synchronised with the video footage to identify each foot as it struck the platform. For each foot stride we measured the maximum and minimum force, as well as the integral of the force-time profile using the trapezoidal numeration integration function (trapz.m) in matlab, for the fore-aft, medial-lateral and dorsal-ventrally directed forces. We corrected for the inclined slope in the fore-aft forces, by multiplying fore-aft force by the cosine of the slope (40 degrees), and removing the sine of the vertical force. Similarly we corrected vertical forces by multiplying it by the sine of the angle and adding the cosine of the fore-aft forces. This rotated forces such that they were comparable to flat-trackway forces.

#### (iv) Excluded strides

To determine the extent to which steady state trials were included in this study we estimated the average acceleration over several strides within the field of view for each run. Acceleration was estimated as the slope of speed over time throughout the run. While runs over the angle and pole surface had a median value around zero, runs over the flat surface show a distinct decelerating trend (Supp Fig. 2). Therefore we excluded all these strides from our main analysis, but we chose to present them in the supplementary material as a means of visual comparison of differences between angled platforms and the pole runs. Caution should be used in the interpretation of the runs along the flat surface, as it will be ambiguous the extent to which changes are the result of surface differences, or result from the inclusion of predominately breaking strides.

#### (v) Statistics

For all analyses, a within subject design was used, including subject as random factor using the lme.R function from the nlme package (Pinheiro et al., 2017) in R (ver 3.2.3 -- “Wooden Christmas-Tree”). To examine variation between factors we specified the model with lme.R function, and then used glht.R function from the multcomp package (Hothorn et al., 2008) to perform Tukey post hoc tests, correcting the P values using the Bonferroni adjustment method.

## Results

### Speed

Quolls with different body lengths did not use significantly different speeds when traversing the two platforms (F_1,10_ = 0.24, P = 0.635). However, quolls ran significantly faster on the wide platform than on the narrow pole (F_1,181_ = 22.8, P < 0.001) (Figure 2).

**Figure 2.**
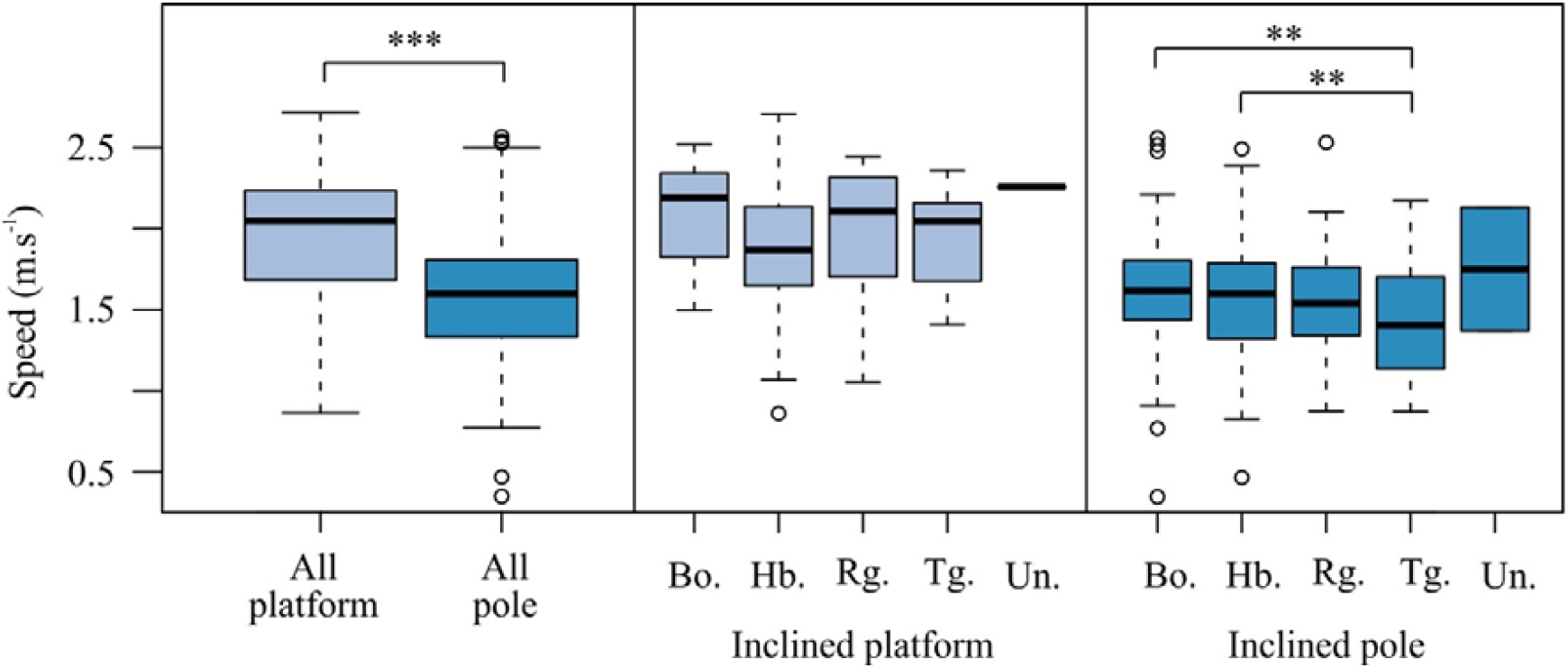
Variation in speed with surface and gait in Northern quolls (*Dasyurus hallucatus).* Speed varies significantly between the surfaces and with gait choice. Bo – Bound, HB – Half bound, Rg – Rotary gallop, Tg – Transverse gallop, Un – Unknown. *** indicates P < 0.001, ** indicates P < 0.01. Boxes represent the median, with hinges representing the first and third quartiles; whiskers represent the 95% CIs, and dots represent outliers.

Speed also varied significantly with gait. For the inclined surfaces, speeds varied between gaits on the narrow pole (F4,206 = 3.68, P = 0.006), but not on the platform (F4,_5_2 = 1.87, P = 0.130). On the narrow pole, transverse gallops were slower than bounds (z = 3.49, P = 0.004), and half bounds (z = 3.44, P = 0.005), but speeds during other gaits were not significantly different. Speed differences with gait were most prominent on the flat platform as reported in supplementary materials.

### Stride characteristics

Duty factor varied with both speed (F_1,266_ = 238.39, P < 0.001) and surface (F_1,266_ = 224.71, P < 0.001), though there was no significant interaction between speed and surface (F_1,266_ = 1.39, P = 0.238). Duty factor was highest on the inclined pole, reflecting the lower speeds on this surface. After removing the interaction term, surface still had a significant effect on the relationship between speed and duty factor, suggesting a significant difference in the intercept between the surfaces (Figure 3 a). Thus, quolls are able to move along the pole at similar speeds to the inclined platform, but do so using a higher duty factor. This increase in duty factor is probably required in order to maintain stability, via increased contact time between the surface and the feet. However, in order to maintain similar speeds at higher duty factors, swing time must be greatly reduced. At any given speed the stride duration tends to be shorter for strides on the inclined pole when compared to the inclined platforms (Surf: F_1,267_ = 6.72, P = 0.010, Speed: F_1,267_ = 291.52, P < 0.001; Fig. 3b).

**Figure 3.**
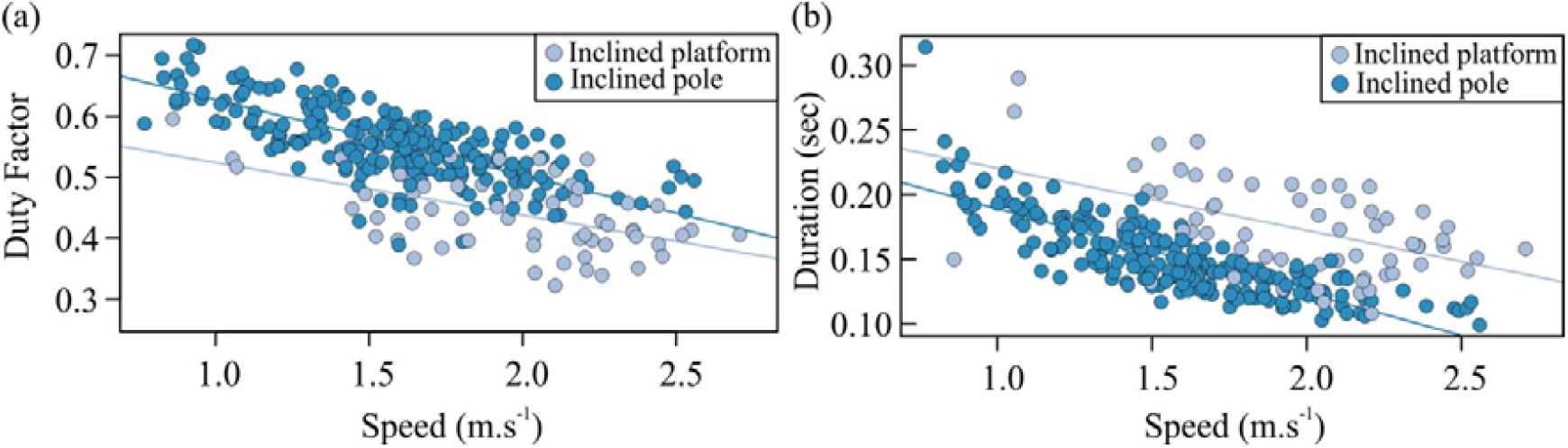
Relationship between speed and duty factor a) and speed and swing time and b) for the inclined platform (light) and inclined pole (dark) conditions. Solid lines indicate linear regression model for each condition.

### Timing between footfalls

Differences in footfall timing between the forefeet (F.lag) was associated with speed, surface, and the interaction between the two (Speed, F_1,263_ = 20.98, P < 0.001; Surface, F_1,263_ = 10.16, P = 0.002; Speed:Surface, F_1,263_ = 4.59, P = 0.033). Removing the interaction term affected the model fit, suggesting significant differences in the slope and intercept between the surfaces (L.ratio = 4.55, P = 0.033). F.lag decreased rapidly with speed on the inclined platform (Fig. 4a) This effect was reduced on the inclined pole, with shorter F.lag being preferred across speeds, indicating the forefeet are acting in unison on narrow poles (Figure 4a). Results from flat surfaces, presented only in the supplementary materials, show a significantly steeper slope than both the inclined platform and the inclined pole (Supp. Figure 5). Overall this suggests that there is a tendency to increase the temporal distance between the forefeet at higher speeds, but this is greatly constrained on narrower surfaces.

**Figure 4.**
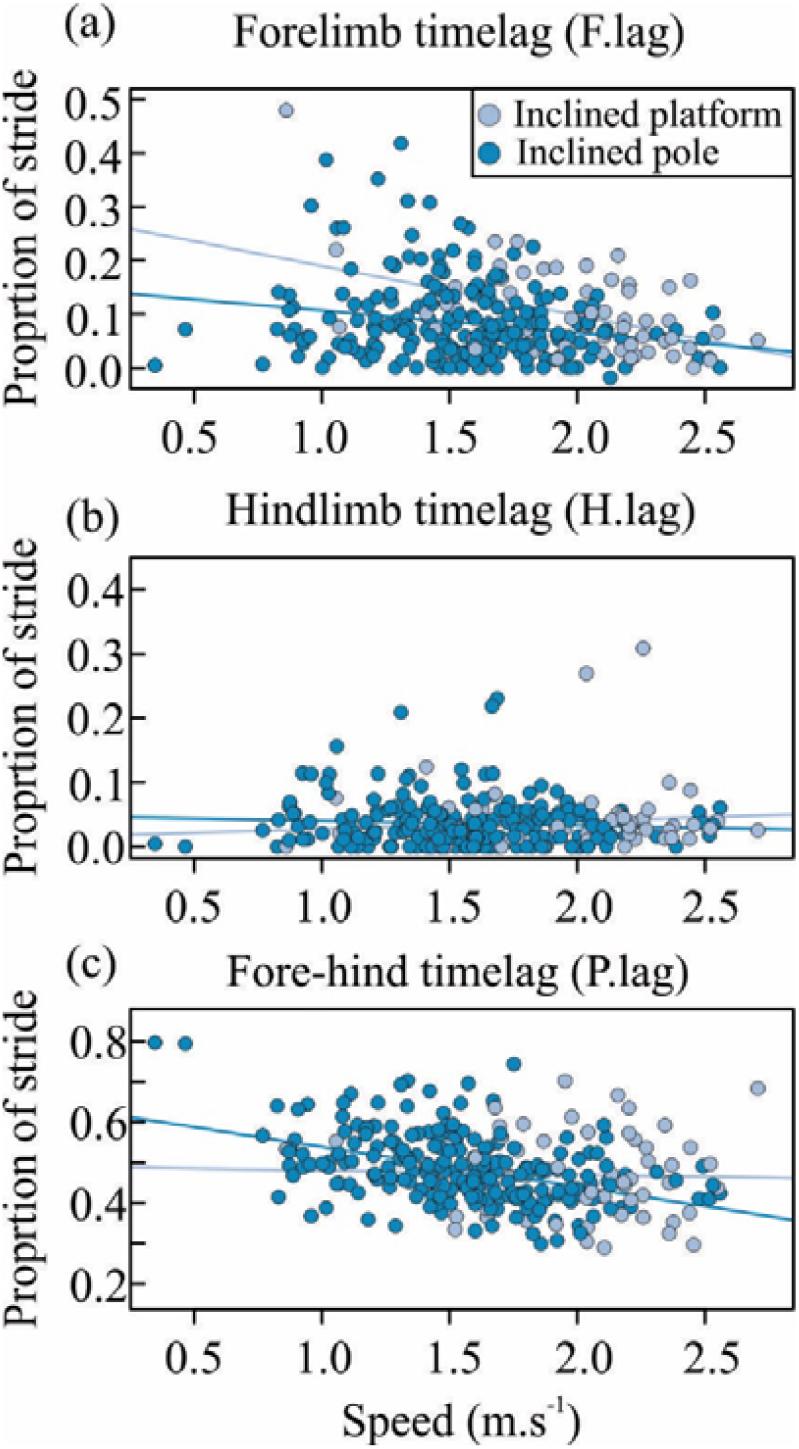
Relationship between a) speed and forelimb timelag (F.lag), b) hindlimb timelag (H.lag) and c) fore-hind timelag (P.lag) for the inclined platform (light) and inclined pole (dark) conditions. Solid lines indicate linear regression model for each condition. F.lag, H.lag, and P.leg were calculated as the temporal lag between the mid-stances of the two forefeet, hindfeet, and averaged across the ipsilateral feet, respectively divided by the total stride time.

Conversely, the time difference between footfalls of the hindfeet (H.lag) was not significantly affected by speed or surface on the inclined surfaces (Square root transformed H.lag: Speed, F_1,266_ = 0.05, P = 0.825; Surface, F_1,266_ = 0.22, P = 0.636; Figure 4b). H.lag was negatively and linearly related to speed for the flat surface (F_1,160_ = 82.2, P < 0.001; See Supplementary Material). On flat surfaces, H.lag tended to be highest for the low speeds, but was greatly restrained with values close to zero (indicating simultaneous footfall of both hindfeet) at higher speeds. Overall hindfeet tend to be used simultaneously (indicated by short time lags) on all speeds and surfaces.

Time lag between the mean forefeet and hindfeet strike (P.lag) was significantly correlated with speed (F_1,266_ = 18.75, P < 0.001; Fig. 4c), but not with surface (F_2,266_ = 1.89, P = 0.169), while the interaction between surface and speed was close to statistical significance (F_1,266_ = 3.63, P = 0.057). When analysing each surface separately, there was no significant effect of speed on P.lag for the inclined platform (F_1,59_ = 0.09, P = 0.758), but a significant effect of speed was found on the inclined pole (F_1,220_ = 34.5, P < 0.001). On this latter surface, higher speeds were associated with a reduction in the phase lag between the fore and hindfeet (Fig. 4c).

### Distance between footfalls

The hindfeet were generally kept much closer together (0.033 ± 0.002 m; mean ± se for all trials) than the forefeet (0.040 ± 0.002 m). The absolute total distance between the forefeet was significantly larger on the inclined platform than the inclined pole (F_1,78_ = 151.17, P < 0.001; Figure 5a), but was not affected by speed (F_1,78_ = 0.98, P = 0.323), nor by the interaction between speed and surface (F_1,78_ = 0.74, P = 0.394). When examining footfall distance in only the fore-aft axis, forelimb distance (F.distX) decreased with speed (F_1,78_ = 6.36, P = 0.014), and was greater on the inclined platform than the inclined pole (F_1,78_ = 8.51, P = 0.005; Figure 5a). Similarly, the interaction between speed and surface did not affect F.distX.

**Figure 5.**
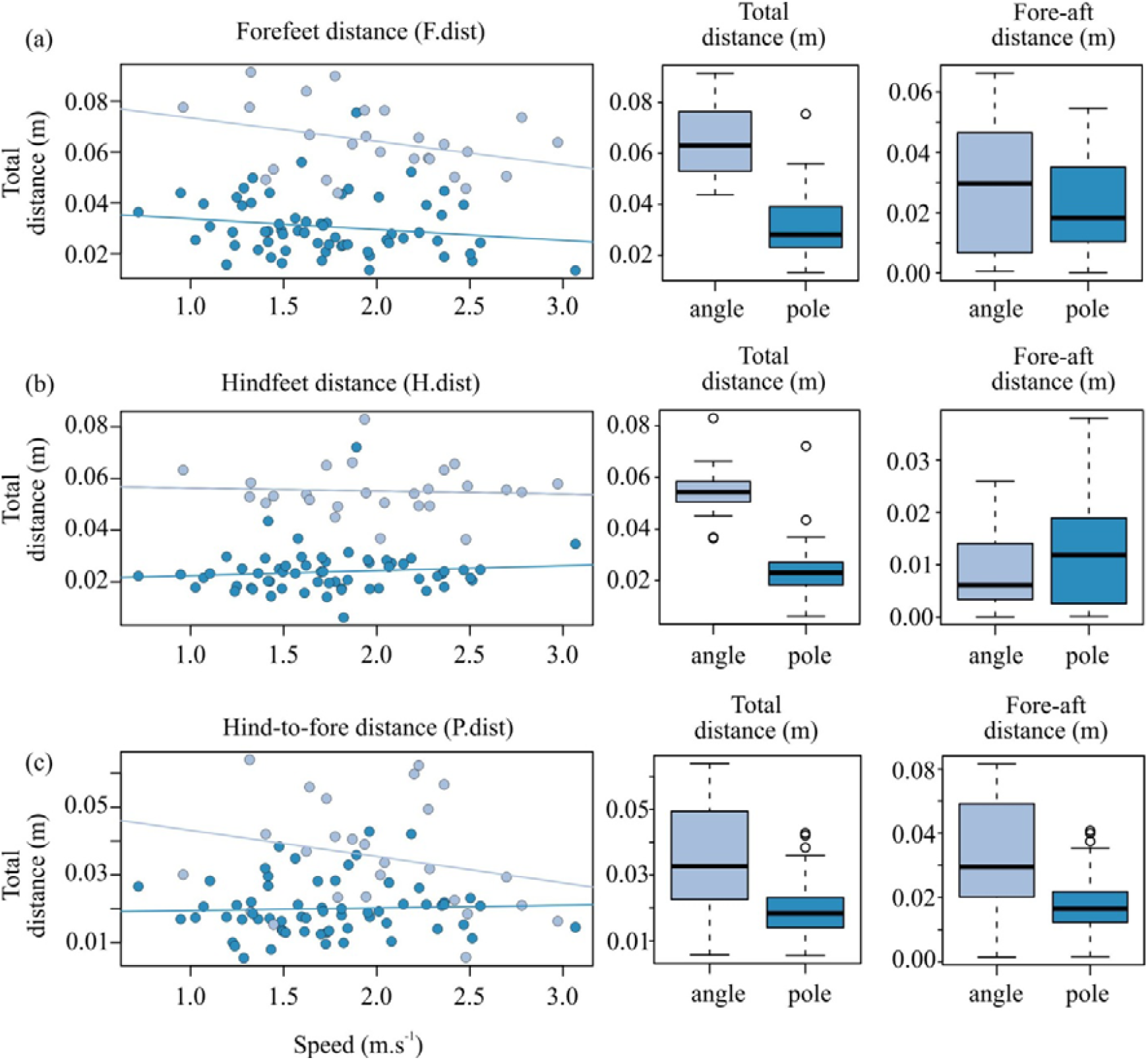
Distances between foot falls; a) for the left and right forefeet (F.dist), b) the left and right hindfeet (H.dist) and c) the mean distance between the forefoot pair and the hindfoot pair (P.dist) for the inclined platform (light) and inclined pole (dark). Given that lateral distances are constrained on the narrow inclined pole, we also report the fore-aft distances between each foot pair.

The absolute distance between the hindfeet was significantly smaller on the inclined pole than the inclined platform (F_1,76_ = 2.05.97, P < 0.001; Fig. 5b). However, unlike the forefeet, the absolute distance between the hindfeet increased significantly with speed (F_1,76_ = 16.12, P < 0.001; Fig. 5a), with no significant interaction between surface and speed (F1,76 = 0.49, P = 0.483). Compared with the absolute distance, the fore-aft distance between the hindfeet shows a considerable drop, below 2 mm and close to the resolution limit of this method (Figure 5b).

Absolute hind-to-fore foot distance was greater on the inclined platform than on the inclined pole (F_1,78_ = 35.73, P < 0.001; Fig. 5c), but was unaffected by speed (F_1,78_ = 0.58, P = 0.446) and the interaction between speed and surface (F_1,78_ = 2.51, P = 0.117). On the flat surface, absolute hind-to-fore foot distance was greater at faster speeds (F_1,54_ = 14.9, P < 0.001), suggesting faster strides could be achieved by reaching further forwards with the forelimb pair (Supplementary material). Yet for both the inclined platform and the inclined pole surfaces, no significant relationship exists between speed and absolute hind-to-fore foot distance. Results for the fore-aft displacement only (P.distX) largely reflect those of P.dist, as might be expected, given that much of the displacement between the fore and hindlimbs is expected to occur along this fore-aft axis.

### Ground reaction forces

To determine the relative contribution of each of the feet towards propulsion, we recorded the output of the forces transferred to the substrate (platform or pole) by each limb individually wherever possible, or by pairs of feet in instances where they touched the force plate simultaneously (Figure 1d).

The integral of the fore-aft force trace with respect to time (impulse) throughout the stance phase was significantly different among the feet (F_5,100_ = 34.3, P < 0.001), but did not significantly differ between the surfaces (F_1,100_ = 1.49, P = 0.224), nor did it vary with speed (F_1,100_ = 0.15, P = 0.698) (Figure 6a). Higher forces were produced by the paired hindfeet compared with the paired forefeet (z = −12.3, P < 0.001), and between the paired hindfeet with either the left (z = −8.0, P < 0.001) or right (z = −6.0, P < 0.001) forefoot alone, with no other comparison being significant. The hindfeet (−0.39 ± 0.026 N.sec) provide over three times the propulsive force of the forelimbs (−0.12 ± 0.010 N.sec) during locomotion on the inclined surfaces. The minimum of the fore-aft force (which represents the peak reaction force against quolls pushing themselves forward) decreased significantly with speed (F_1,100_ = 9.43, P = 0.003), indicating an increase in fore-aft force with increasing locomotor speed.

**Figure 6.**
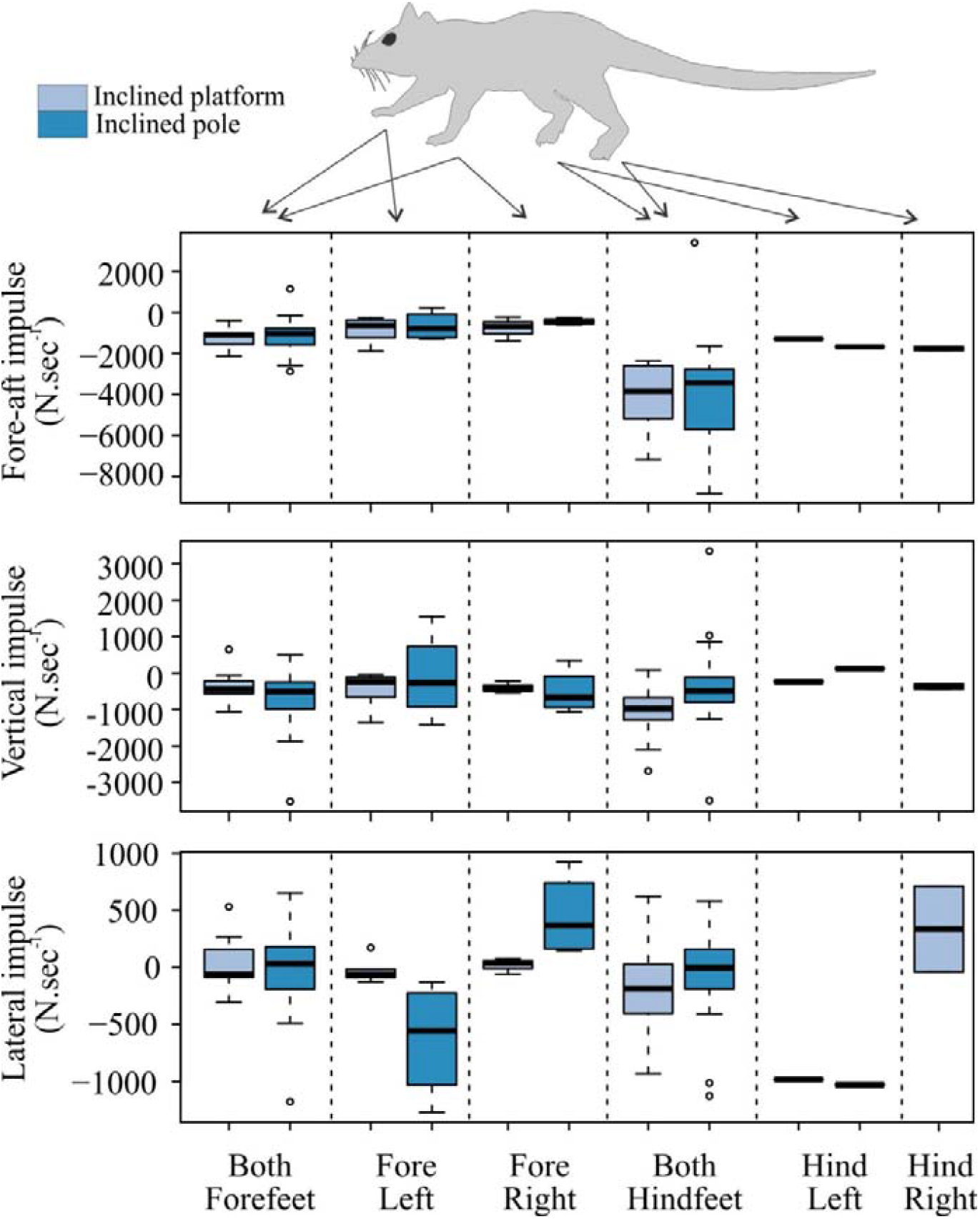
Impulse determining the relative contribution of each of the feet where possible, or for pairs of feet where they touch the force plate simultaneously for the Northern quoll (*Dasyurus hallucatus).* Impulse was calculated as the integral of the force trace with respect to time for a) the fore-aft, b) vertical, and c) lateral directions for inclined platform (light) and inclined pole (dark). Positive lateral impulses indicate force directed towards the left and vice versa. Box and whisker plots as per Figure 2.

Like impulse, there was no effect of surface (F_1,100_ = 0.41, P = 0.521) on minimum fore-aft force, but there was significant variation among feet (F_5,100_ = 52.2, P < 0.001). As for the impulse above, the hindfeet provided the greatest propulsive force.

The integral of the vertical force trace throughout the stance phase was not affected by surface (F_1,100_ = 1.08, P = 0.302), foot (F_1,100_ = 0.67, P = 0.647), nor speed (F_1,100_ = 0.00, P = 0.996), suggesting that all feet contribute near equally to body weight support during movement (Figure 6b). The mean vertical impulse for all feet was −0.055 ± 0.0072 N.sec. The minimum of the vertical force (which represents the peak reaction force against quoll body weight) was not affected by speed (F_1,100_ = 0.11, P = 0.743) or surface (F_1,100_ = 0.01, P = 0.935), but was weakly affected by foot (F_5,100_ = 2.78, P = 0.022), with a post-hoc test showing only differences between the fore-left foot alone, with the combined forefeet (z = 3.15, P = 0.024)

The maximum vertical force, which indicates whether the surface is pulled toward the body during the stance phase, decreased significantly with speed (F_1,100_ = 4.97, P = 0.028), with slower strides requiring a greater pulling force. Maximum vertical force was greategr on the inclined pole than the inclined platform (1.07 ± 0.155 N vs 0.36 ± 0.112 N; F_1,100_ = 4.15, P = 0.044), likely as a result of quolls being better able to grip underneath the narrow pole. However, maximum vertical force was not significantly different between feet (F_1,100_ = 0.55, P = 0.732).

The lateral force trace integral was not significantly affected by speed (F_1,100_ = 0.24, P = 0.626) or surface (F_1,100_ = 0.01, P = 0.937), but like the vertical and fore-aft impulse, there was a significant effect among feet (F_5,100_ = 5.91, P < 0.001) (Figure 6c). Post-hoc tests highlighted that the single forelimbs (fore-left and fore-right) were capable of producing significantly different lateral forces, yet this trend was only evident on the narrow pole (Figure 6c).

### Corrective Torques

We explored the magnitude of corrective torques applied along the narrow inclined pole (Fig. 7). We do not report these for the inclined platform, as they could unlikely contribute to stability on a flat surface, where forelimbs cannot grasp. Neither counter-clockwise nor clockwise torque was significantly related to speed or foot (counter-clockwise, speed: F_1,59_ = 0.37, P = 0.543, foot: F_4,59_ = 1.14, P = 0.348; clockwise, speed: F_1,59_ = 0.02, P = 0.879, foot: F_4,59_ = 1.57, P = 0.193). However, when comparing simultaneous foot falls, the paired forefeet and the paired hindfeet produced substantial corrective torques, likely related with the decreased distance between fore and hind feet pairs on the inclined pole in comparison to the inclined platform. The left forelimb produced larger clockwise corrective torques in comparison to the right forelimb (FL = 72.15±18.96 N.mm, FR = 16.22±5.41 N.mm; n=4). In contrast, the right forelimb produced greater counter-clockwise corrective torques than the left forelimb (FL = −16.98±14.17 N.mm, FR = −51.64±16.83 N.mm; n=4). The direction of these corrective torques is consistent with each single forelimb being used in a gripping while pulling forefoot stride.

**Figure 7.**
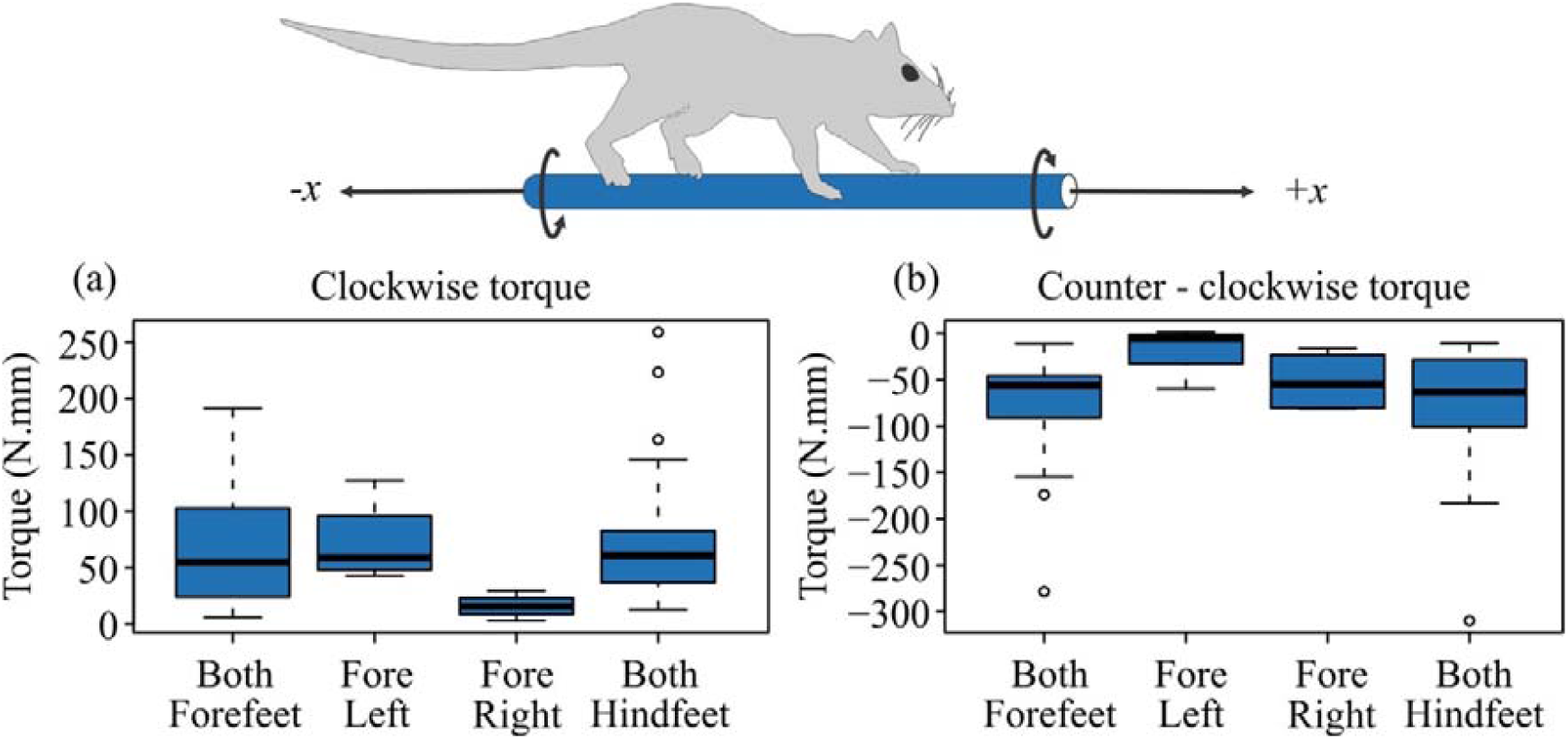
Maximum corrective torques in the a) clockwise and b) counter clockwise directions on the inclined pole for the different combinations of foot limb pairs for the Northern quoll (*Dasyurus hallucatus).* The direction of torques is consistent with the right-hand rule. Sufficient data was not available for single hindlimbs. Box and whisker plots as per Figure 2.

## Discussion

Understanding the extent to which a species has adapted to a specific environment often can require the combination of relevant performance measures and knowledge of the biomechanical limitations imposed by that environment. Greater declines in populations of Northern quoll from open grasslands compared with semi-arboreal rocky habitats suggest greater predation pressure in open habitats. This presents a unique system to quantify the link between performance and survival in natural habitats. To explore the extent to which habitat can compromise the performance of Northern quolls (*Dasyurus hallucatus*) we analysed their kinematics and kinetics while they moved along terrains designed to represent aspects of their natural environment.

### Environment-dependent locomotion in the Northern quoll

We found that quolls used slower speeds when moving on a narrower surface, but used similar gaits on both wide and narrow inclined structures. This suggests that a quolls’ gait choice is independent of both speed and the structure of the terrain. This highlights the limitations of examining only variation in classical gait characterizations when comparing the movement of small, agile animals within habitats of differing complexity. Therefore, we also measured variation in the kinematic and kinetic patterns of quoll movement.

Quolls were capable of moving at similar speeds along the inclined pole as the inclined platform, but did so using a much higher duty factor. Duty factor is the percentage of the stride cycle when the foot is in contact with the substrate, suggesting relatively longer stance phases may be required to provide stabilizing moments on narrow substrates, such as the inclined pole. In order to maintain similar speeds when using higher duty factors, the quolls swung their legs faster on the inclined pole. This may provide evidence for an alternative neuromuscular strategy for arboreal movement in quolls that may not be available to other species. This increased leg swing speed may be responsible for the increase in the probability of mistakes, which is present when quolls run at higher speeds, or on narrower surface (Nasir et al., 2017).

We found a similar pattern in the distance and timing of footfalls, whereby distance and time both decreased at faster speeds and on narrower surfaces. This too suggests a change in the biomechanical strategy employed by quolls on narrow surfaces, reflecting a coupled use of limb pairs at higher speeds. Coupled limb behaviour is likely associated with the corrective forces and torques necessary to maintain stability on narrow surfaces (Cartmill, 1985). We found greater changes in both lateral forces and the ability to produce corrective torques on narrow, as compared to wide surfaces. Surprisingly, quolls were not only able to produce corrective torques with simultaneous limb pairs, as previously shown in primates (Chadwell and Young, 2015), but also with single limb surface contacts. These former corrective torques, between left and right limbs of a girdle, likely explains why we observed a decrease in the timing and distance of footfalls, suggesting that like primates, quolls expand their effective grasp by gripping the substrate between limb pairs. But this latter single limb torque pattern indicates a level of complexity in the forefeet which appear in some cases to grip the narrow surface using a single foot and produce significant corrective ‘pulling’ torques.

### Cats and dogs: changes in gait parameters

The decreased population density of Northern quolls in open grassland habitats is often attributed to introduced predators such as cats and dogs, and a decrease in habitat complexity due to land clearing, altered fire regimes, or grazing by invasive herbivores (Newsome, 1975, Burbidge and McKenzie, 1989, McKenzie et al., 2007, Oakwood, 2000). A comparison of the biomechanical capacities between quolls, cats, and dogs may therefore provide insight into the relative performance of these predators and quolls in different habitats. Galves-Lopez *et al.* (Gálvez-López et al., 2011) compared the kinematics of cats and dogs on a raised narrow wooden beam to that of unconstrained overground locomotion, mirroring the experimental protocol used in this study. Similar to quolls, cats adopted lower speeds on narrow supports as compared to overground surfaces, but unlike quolls, swing phase duration was independent of speed during narrow support locomotion (Gálvez-López et al., 2011). During overground locomotion, the swing phase duration of cats decreased with increasing speed, but did not change on narrow surfaces, suggesting that cats are unable to take faster steps on narrow structures but instead appear to take longer strides. In contrast, quolls use an absolutely lower swing phase duration at any speed on narrow structures, implying different control strategies between cats and quolls. Further, unlike quolls, where we found a significant decrease in the timing between forefeet contact among narrow and wide surfaces, cats deviate from this pattern and increase the timing between forefeet contacts on these narrow structures. The extent to which this limits their ability to produce corrective torques, as found in quolls, remains to be shown.

In contrast to quolls and cats, which both decrease their speeds on narrower surfaces, dogs adopt higher speeds. These higher speeds are achieved via an increase in both stride frequency and stride length, and a decrease in stance phase duration, leading to lower duty factors on narrower substrates (Gálvez-López et al., 2011). Combined, the biomechanical traits evident in dogs on narrow substrates suggest a reduced degree of static stability and an increased reliance on dynamic stability. Static stability is the process by which an animal remains stable because the forces and torques produced by gravity are near equal and opposite to the ground reaction forces and moments (Lammers and Zurcher, 2011a).

However, at higher speeds it becomes increasingly difficult to maintain static stability. Dynamic stability is the process whereby an animal remains stable due to the presence of an angular momentum created by the cyclical motion of the limbs. This cyclical motion produces a ‘gyroscopic effect’ where if a small disturbance changes the lean of a body, the gravitational force produces a restoring torque about the fore-aft axis to maintain stability. Truly arboreal species appear to avoid highly dynamically stabile gaits (Lammers and Zurcher, 2011a), which is likely related to the discontinuous, multi-dimensional, and frequently unstable characteristics of arboreal habitats. This suggests larger terrestrial animals, like dogs, may be at risk of increasing the probability of mistakes on narrow substrates, though this remains to be shown. However, this may not be true in small arboreal specialists, like chipmunks who exploit dynamic stability during locomotion (Lammers and Zurcher, 2011b).

### How do Northern quolls compare with other arboreal species?

Kinematic comparisons of semi-aboreal Northern quolls with other arboreal species may help to determine the extent to which quolls are selected for an arboreal versus terrestrial environment. Like quolls, among 7 species of marsupials (opossums), relative velocities increased with support diameter (Delciellos and Vieira, 2009), indicating a transition to slower and more stable gaits on narrower structures. Similar control strategies to quolls were evident among mouse lemurs, where limb contact time increased on narrow structures, likely to aid stability through an increase in the time available to apply corrective torques (Shapiro et al., 2016). Though this latter species did differ by showing an increase (rather than a decrease in quolls) in the time between the trailing and leading limbs (P.lag) on narrow surfaces.

Greater differences were observed between quolls and rats (Camargo et al., 2015). Arboreal rat species tend to increase stride frequency and decrease stride length on narrow structures, compared to terrestrial species that decrease both stride length and frequency. Thus, arboreal species show an increase in speed on narrow structures, whereas terrestrial species slow down (Camargo et al., 2015). The biomechanical control strategies used by quolls is therefore more akin to terrestrial than arboreal rats.

A kinematic pattern common among arboreal specialists, particularly among primates, is the use of a diagonal-sequence footfall pattern during walking, where the fore-left and hind-right limbs (or vice-versa) are used in conjunction during stance (Muybridge, 1887, Prost, 1965, Prost, 1969, Hildebrand, 1966, Hildebrand, 1977, Hildebrand, 1980, Hildebrand, 1985, Vilensky and Larson, 1989, Lemelin and Grafton, 1998, Cartmill et al., 2002). This footfall pattern is often thought to be beneficial when used in association with grasping extremities when moving and foraging on thin flexible branches. This theory was tested among the grey short-tailed opossum (*Monodelphis domestica*) and the woolly opossum (*Caluromys philander*) – and showed that the latter species, which possessed more developed grasping extremities, showed a greater use of diagonal-sequence walking gaits (Lemelin et al., 2003). In the current study, this walking sequence was not observed among quolls, with P.lag values approximating 0.5, indicating forefeet or hindfeet pairs are used (rather than diagonals) during bounds and half-bounds. Thus despite at least some ability to grasp, quolls display gait characteristics which suggest an important role of terrestrial locomotion and an increased reliance on dynamic stability while on narrow structures.

Kinetic comparisons also showed differences between classically arboreal species and quolls. Like quolls, the grey short-tailed opossum (*Monodelphis domestica*) showed both fore- and hind limbs had equal roles in body weight support (vertical force) on inclined surfaces (Lammers and Biknevicius, 2004). However, differences arose in propulsive forces (fore-aft): in quolls, the hindlimbs produced the majority of the propulsive impulse whereas in opossums, it was the forelimbs which exerted the greater propulsive impulse (Lammers et al., 2006).

It has further been hypothesised that much of the kinematic changes to gait on narrow supports are linked to a functional reduction in the vertical impulse, which would reduce oscillations of the centre of mass and increase stability on narrow structures (Gálvez-López et al., 2011, Schmitt and Lemelin, 2002, Schmitt et al., 2006, Young, 2009, Lemelin and Cartmill, 2010). However, in the current study we found little or no evidence for a decrease in vertical impulse on narrow supports which may function to reduce these oscillations. In other species, vertical impulses were lower on arboreal supports for the common marmoset (*Callithrix jacchus*) (Chadwell and Young, 2015) and the grey short-tailed opossum (Lammers and Biknevicius, 2004), although the authors caution that in this latter species, this may be attributed to differences in speed between the treatments.

The torques generated by the limbs about the long axis of a branch during locomotion may also be important for stable locomotion on arboreal substrates. Although several characteristics of quoll locomotion resemble terrestrial species, quolls appear more similar to arboreal species in their ability to produce stabilizing torques on narrow supports. For example, the grey short-tailed opossum (*Monodelphis domestica*) uses long-axis torque to avoid toppling on a branch, though in this species the forelimbs generated significantly greater torque than the hindlimbs which is probably explained by the greater weight bearing role of the former (Lammers and Gauntner, 2008). Similarly, the Siberian chipmunk (*Tamias sibiricus*) produced considerable torque to counteract rolling moments which were equal between the fore and hindlimbs (Lammers and Zurcher, 2011b). Further, like quolls, the total impulse of the rolling torques in these chipmunks was usually greater than or less than zero (ie: not balanced within a stride). This indicates that like chipmunks, quolls may balance out the torques acting on the centre of mass over the course of two or more strides to maintain stability.

### A biomechanical framework to predict the survival of key species

These comparisons suggest that there are multiple biomechanical strategies available to achieve support on narrow substrates, and the degree to which different species exploit these strategies varies along a spectrum from fully terrestrial, to semi-arboreal, to fully arboreal. Northern quolls are often described as semi-arboreal specialists (Schmitt et al., 1989). Consistent with this we show many of the gait characteristics associated with quolls appear similar with terrestrial species, with some characteristics useful in arboreal habitats.

Historically Northern quolls occupied both floodplains (terrestrial) and rocky (semi-arboreal) habitats. Our biomechanical analyses suggest that cats and dogs are likely superior in terrestrial environments, owing to their larger body size and greater speeds (Gálvez-López et al., 2011). This may partially explain the declines and/or reduced populations of Northern quolls in these terrestrial environments (Oakwood, 2000). Conversely, quolls appear to have biomechanical characteristics consistent with a stability advantage at higher speeds on narrow supports when compared to cats and dogs (Gálvez-López et al., 2011), likely explaining why the decrease in quoll populations in semi-arboreal habitats (such as rocky outcrops) has not been as severe (Oakwood, 2000). However, quolls did not show many of the characteristics common in truly arboreal specialists (e.g. diagonal sequence gaits) such as opossums, which may explain why populations of quolls are showing more obvious signs of decline than other native species such as possums (Woinarski et al., 2001). Quolls appear to share a larger percentage of their performance space with cats and dogs, and this overlap, combined with reductions in habitat complexity, may be directly related to their declining population on mainland Australia.

Here we show the value in using biomechanical analyses to predict the survival and fitness of key species within a habitat (Wilson et al., 2018). The relative performance of a species, results from its ability to employ biomechanical strategies necessary to maintain stability and outperform key predators (invasive or otherwise) within that habitat. This concept may become even more important as we move to a period of increased human impact within the Anthropocene, and may allow us to predict the influence of climate change, urbanization, and deforestation on species of conservation importance.

## Acknowledgements

We thank members and volunteers of the Wilson Performance Lab for assistance with running the experiments in the field. We also thank the Anindilyakwa Land and Sea Rangers of Groote Eylandt for their generous assistance, logistical support and use of laboratory facilities. We also thank the traditional owners of Groote Eylandt for their generous support and access to their land.

